# Temporal and spatial variability in availability bias has consequences for marine bird abundance estimates during the non-breeding season

**DOI:** 10.1101/2024.03.13.584773

**Authors:** Ruth E Dunn, James Duckworth, Susan O’Brien, Robert W Furness, Lila Buckingham, Francis Daunt, Maria Bogdanova, Jonathan A. Green

## Abstract

1. To effectively monitor how marine ecosystems are being reshaped by anthropogenic pressures, we require understanding of species abundances and distributions. Due to their socio-economic and ecological value, predatory species are often at the forefront of survey efforts. However, survey data are only valuable if they can reliably be converted into estimates of underlying distributions.
2. We consider at-sea surveys of marine predators that often inform ecological impact assessments of offshore windfarms. These surveys are subject to a form of detection bias called ‘availability bias’ whereby individuals which are submerged below the surface are consequently ‘unavailable’ for detection. Although correction factors are commonly used in these surveys, they are currently based on limited data that may not be species-, time-, or area-specific. Here, we use time-depth-recorder data to investigate variation in marine bird availability bias.
3. We found that the proportion of diving marine birds submerged below the sea surface during daylight hours, and therefore unavailable to be counted during surveys, varied by species, month, and area. For three of our focal species wintering around northwest Europe (Atlantic puffin, common guillemot, razorbill) our results were different to comparable values previously used to correct for the availability bias, whereas no correction factors are regularly used for the fourth species (red-throated diver). We now present availability bias correction factors that are species- and month-specific to the areas the study populations use during their non-breeding seasons: the North Sea, the north and west coasts of the UK, the Baltic Sea, and Icelandic coastal waters.
4. *Synthesis and applications:* Variation in the proportion of daylight hours that marine birds spent submerged lead to differences in availability bias correction factors, thereby impacting estimations of their abundances. We encourage use of correction factors that use data from the species, marine area, and month during which surveys are conducted to provide more accurate abundance estimates. Using more relevant correction factors will result in increasingly accurate abundance and distribution estimates of diving marine birds, with relevance for a range of applications including planning for offshore windfarm developments, the designation and monitoring of protected areas, and understanding environmental change.

## Introduction

Anthropogenic pressures are rapidly reshaping terrestrial, aquatic, and marine ecosystems (Delong et al., 2018; Henson et al., 2017). As we navigate the challenges of the Anthropocene, strive to monitor and manage changes to these different ecosystems effectively, and reduce further harm, an understanding of species abundances and distributions, and how these are changing, is essential (Boivin et al., 2016; Halpern et al., 2015). Examples of this include when researching and documenting the influence of environmental change on species distributions (Jetz et al., 2019), the consideration and monitoring of protected areas (Edgar et al., 2014), and the execution of environmental impact assessments (EIAs) for oil and gas licensing and offshore windfarms (Dierschke et al., 2016). However, if methods used to establish species abundances and distributions are flawed, then this is likely to lead to poor decision making and perhaps even exacerbate anthropogenic environmental problems (Guisan et al., 2013).

When devising management and conservation plans, apex predators tend to be highly regarded due to their social, economic, and ecological value (Sergio et al., 2006). Despite their significance, estimating the abundances and distributions of marine predators is challenging. This is an issue of high importance, since, in an effort to improve energy security and reduce human reliance on fossil fuels, there has been expansion in the development of marine renewable energy technologies that harness natural resources in the habitats occupied by these species (Soares-Ramos et al., 2020). As part of offshore windfarm EIAs, the at-sea abundances and distributions of marine apex predators are typically assessed using either visual observations, from ships or aircrafts, or digital aerial surveys (Buckland et al., 2012; Thaxter & Burton, 2009). Whilst ideally these surveys would capture and record the presence of all individuals, many taxa, including marine reptiles, mammals, and birds, spend extended periods below the water’s surface during which they are undetectable by the methods used to survey them. Historically, this detection bias (sometimes referred to as ‘visibility bias’, but hereafter referred to as ‘availability bias’) has been corrected for in these taxa by following methods derived to estimate densities of harbour porpoises *Phocoena phocoena* (Barlow et al., 1988), despite the fact that marine birds can engage in additional relevant behaviours, most notably flight. Thus whilst the quantification of at-sea marine bird abundances and distributions is an important issue, it is also imperative that the approaches used to derive correction factors that account for availability bias are appropriate for this taxon (Certain & Bretagnolle, 2008; Winiarski et al., 2014).

Diving marine birds such as Atlantic puffins *Fratercula arctica* (hereafter ‘puffins’), common guillemots *Uria aalge* (hereafter ‘guillemots’), razorbills *Alca torda*, and red-throated divers *Gavia stellata* are potentially sensitive to industrial activities, including offshore windfarm developments, across their ranges (Dierschke et al., 2016). Accurately surveying the abundances and distributions of these species is therefore key and historical estimates of at-sea marine birds were likely to have been substantial underestimations, illustrated via mismatches in at-sea survey estimates and predictions of wintering population derived from colony counts (Harris & Wanless, 2011). Over the last decade, offshore windfarm EIAs have attempted to account for availability bias, by considering the time that individuals from particular locations at specific times of year spend submerged, as a proportion of total time spent on water. At present, correction factors are applied only to birds visible on the water’s surface during surveys, with birds in flight being added later (e.g., Harker et al., 2022). Often only a single correction factor is used for each species, this factor frequently having been measured during the breeding season and then applied over the whole annual cycle. For example, when accounting for availability bias within monthly at-sea digital aerial surveys of marine birds in waters off south-east Scotland across an annual cycle, Harker et al. (2022) followed other similar assessments in using values for puffins derived from those breeding at Petit Manan Island, Maine, USA (Spencer, 2012), for guillemots and razorbills from those breeding at the Isle of May, Scotland (Thaxter et al., 2010), and for red-throated divers, no correction factors are regularly used (Irwin et al., 2019). However, the ever increasing number of biologging studies from across ocean basins (e.g. Lescure et al., 2023), means that there is no longer a need to rely on these commonly used factors from specific areas. Indeed, continued advances in biologging provide opportunities to quantify the time that different species of diving marine birds spend above or below the surface of the sea at different times of year (Duckworth et al., 2021), enabling the derivation of correction factors for availability bias throughout the annual cycle. Together, this increased understanding now enables us to fine tune marine bird abundance estimates, accounting for variation in behaviour in both space and time.

Marine bird diving behaviour varies spatially, temporally, and interspecifically, being influenced by foraging strategies, energy requirements, life cycle stages, and environmental conditions (Phillips et al., 2017). Puffins, guillemots, razorbills, and red-throated divers breed once annually during the late spring – summer at sites around the coastlines of the Atlantic and Pacific oceans, flying and diving intensively during this period as they forage both to feed themselves and their chicks (Dunn et al., 2019; Eriksson et al., 1990). These species then spend the rest of the annual cycle (the non-breeding season) largely at sea, flying less and often experiencing harsher weather conditions and heightened energetic costs, with consequences for their diving behaviour that vary between species (Duckworth et al., 2021; Dunn et al., 2019, 2020). Furthermore, whilst these marine birds are highly adapted to foraging on benthic and pelagic prey (Kleinschmidt et al., 2019; Linnebjerg et al., 2013), they differ in their ability to forage uniformly throughout the diel cycle, with the diving behaviour of puffins and red-throated divers, in particular, largely being limited to daylight hours (Duckworth et al., 2020; Shoji et al., 2015). Throughout the autumn and winter, as daylight hours become increasingly constrained, varying latitudinally and therefore between non-breeding areas, the proportion of daylight hours that these birds spend submerged may increase. Variation in behaviour over time and between populations, caused by these various drivers, will therefore result in differing magnitudes of availability bias. Failing to account for this variation will lead to inaccuracies in estimates of marine bird abundance.

Here, we use published biologging data on the dive behaviour of puffins, guillemots, razorbills, and red-throated divers wintering across northwest Europe to quantify broad scale temporal and spatial variation in the proportion of time that they spend below the water’s surface. From these data, we calculate new species-, month- and area-specific correction factors for puffins and razorbills in the North Sea, guillemots in the North Sea or off the west coast of the UK, and red-throated divers in the North Sea and Baltic Sea, within eastern Icelandic coastal waters, and around the coast of northern Scotland in the months following their breeding seasons. With these examples, we show the importance of accounting for temporal and spatial differences in availability bias.

## Materials and methods

### Biologging deployments

Breeding adult puffins, guillemots, razorbills, and red-throated divers were captured from their breeding sites at sites in Scotland, Finland, and Iceland (Figure 1) during the breeding season (for details, see Table 1). All birds were fitted with time-depth-recorders (G5, CEFAS, Lowestoft UK, 31 × 8 mm, 2.7 g in air) attached to Darvic leg-rings. In all cases the attachment process took <10 min and the weight of the logger was <2% of the total body mass of the bird. Birds were recaptured during subsequent breeding seasons and the devices were removed. For more details on the puffin and razorbill deployments see Dunn et al. (2019), for guillemot deployments see Buckingham et al. (2023), and for red-throated diver deployments see Duckworth et al. (2021). Sampling periods varied (Table 1) due to some TDRs failing during the autumn and winter.

**Figure 1.**
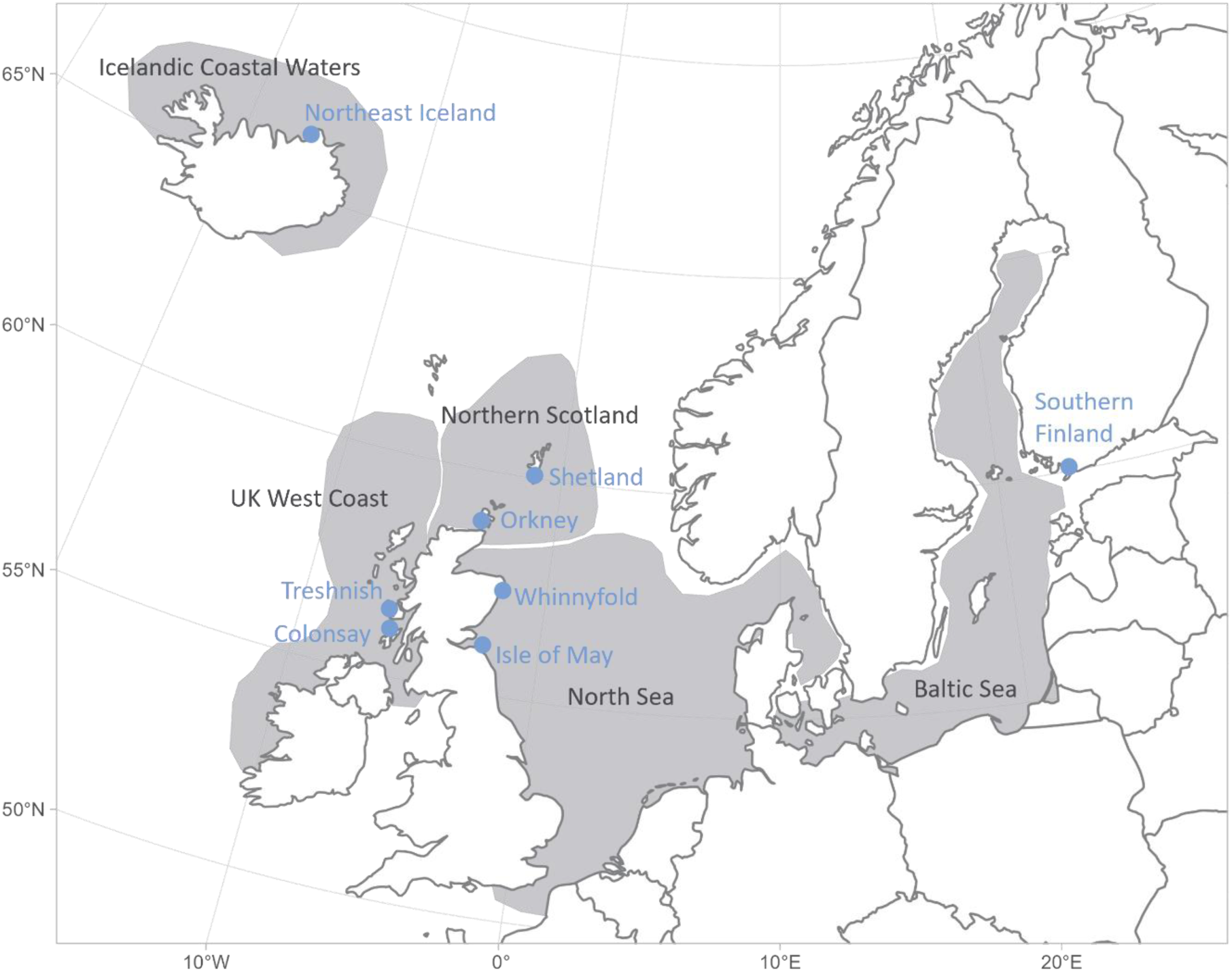
Schematic map illustrating the locations of the breeding sites (shown as blue points and labelled with blue text) where time-depth-recorder loggers were deployed on and retrieved from Atlantic puffins, common guillemots, razorbills, and red-throated divers. Wintering areas are shaded in grey and labelled with black text and are adapted from (Buckingham et al., 2023; Duckworth et al., 2022; Harris et al., 2010). Table 1 details the wintering areas used by each population.

**Table 1.**
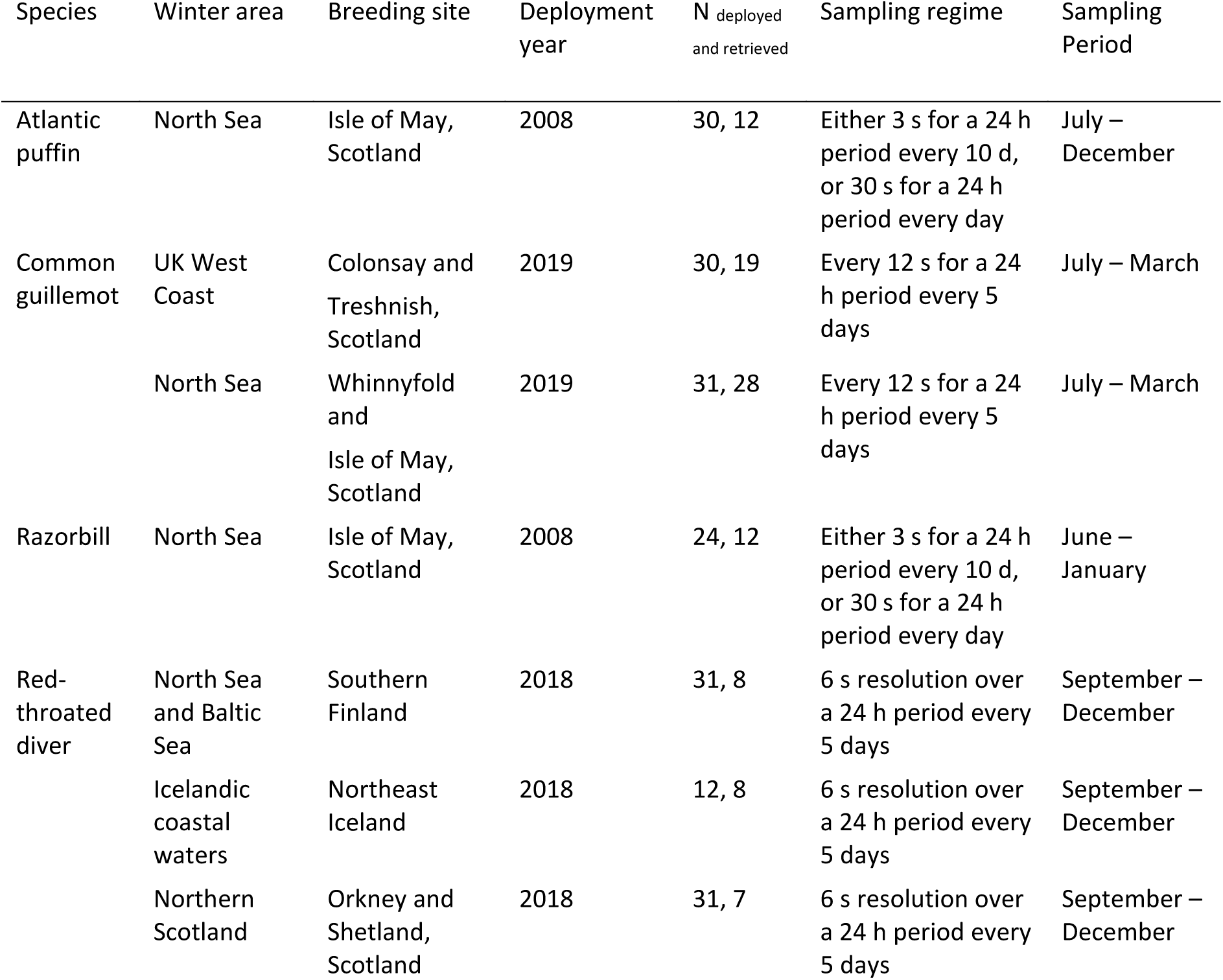
Details of the time-depth-recorder loggers deployed on and retrieved from Atlantic puffins, common guillemots, razorbills, and red-throated divers at breeding sites across northwest Europe. For more details on the puffin and razorbill deployments see (Dunn et al., 2019), for guillemot deployments see (Buckingham et al., 2023), and for red-throated diver deployments see (Duckworth et al., 2021).

### Data processing

All data processing and analyses were conducted in R version 4.2.2 (R Core Team, 2022). First, we curtailed our analyses to the non-breeding period by removing data gathered prior to mean population fledging dates (Duckworth et al., 2022; Dunn et al., 2019; Wanless et al., 2023). We then extracted all incidences where time-depth-recorder depth values exceeded 1 m. To categorise these incidences of submergence behaviour as occurring during either daylight, twilight, or nighttime, we extracted time- and population-specific area estimates. To do this, we obtained centroids of each population’s monthly non-breeding distribution from previous work using light-based geolocator devices (Buckingham et al., 2022, 2023; Duckworth et al., 2022; Harris et al., 2010) (Supporting Information S1). We used the ‘*oce*’ R package to extract sun elevation angles at these areas and categorised each submergence incidence as occurring during daylight (sun was >0 degrees above the horizon), twilight (sun was 0–15 degrees below the horizon), or nighttime (sun was >15 degrees below the horizon) accordingly. We used a cut-off of 15 degrees below the horizon to account for uncertainty in our area estimates. We used the same approach to estimate the duration of daylight hours for each day. We conducted a sensitivity analysis to investigate the impact of using the population’s monthly centroid on our results using the guillemot data (due to them having the most data available and a large latitudinal spread). We calculated the mean latitude plus and minus the standard deviation from all the locations for each population in each month, and this showed that using the lower and upper standard deviation values (instead of the centroid) led to negligible differences in our results (Supporting Information S2).

We counted all incidences of submergence that took place during daylight hours, multiplied this by the sampling interval for that individual, and divided this value by the duration of daylight to calculate the proportion of daylight hours during which each individual was submerged. We excluded twilight and nighttime hours from our analyses as visual marine bird surveys occur during daylight hours only (NatureScot, 2023b). This metric encompasses all behaviour and should be applied to the total count of birds typically captured during daytime aerial surveys, including both those on the surface and in flight. By doing this, we assumed that the proportions of time that birds spend engaged in their key behaviours (i.e., flying, diving and on water) are equivalent to the proportion of individual birds counted (or not, when they are submerged) engaging in these behaviours during surveys. This assumption underpins similar corrections of survey data which are made using time-budget data (Barlow et al., 1988).

### Statistical analyses

We composed a set of generalised linear mixed effects models to quantify the proportion of daylight hours spent submerged by each species based on the explanatory terms available for that species, using the Stan programming language via the ‘*brms*’ R package (Bürkner, 2017). As we were modelling proportional data, we used the Beta distribution that provided flexibility but was limited to between 0 and 1 (Heiss, 2021). For the razorbill and puffin models we included an explanatory term of month only (because data were only available for one area; Tables 1 & 2), whereas for the red-throated diver and guillemot models we include month, area, and their interaction (Table 2). We included month as a categorical representation of time within our models, rather than as a continuous covariate, because marine bird surveys are typically conducted at monthly intervals (NatureScot, 2023a). For all models we included individual bird ID as a random effect.

**Table 2.** Details of the generalised linear models run to model the proportion of daylight hours spent submerged by Atlantic puffins, common guillemots, razorbills, and red-throated divers from breeding sites across northwest Europe during the non-breeding season.

| Model | Explanatory variables |  | Categories | Intercept prior |
| --- | --- | --- | --- | --- |
| Puffin | Single variable: | Month | Aug, Sep, Oct, Nov | $N(-1.802, 0.1)$ |
| Guillemot | Two-way interaction: | Month | Jul, Aug, Sep, Oct, Nov, Dec, Jan, Feb, Mar | $N(-1.150, 0.1)$ |
|  |  | Area | UK East Coast, UK West Coast |  |
| Razorbill | Single variable: | Month | Jul, Aug, Sep, Oct, Nov, Dec, Jan | $N(-1.504, 0.1)$ |
| Red-throated diver | Two-way interaction: | Month | Sep, Oct, Nov, Dec, Jan, Feb, Mar | $N(-0.847, 0.1)$ |
|  |  | Area | Baltic Sea and North Sea, Icelandic coastal waters, Northern Scotland |  |

We adjusted the priors of the coefficients so that they had a normal distribution centred at 0 with a standard deviation of 1 and kept the default brms random effect priors. For the model intercepts, we incorporated prior knowledge in the form of values that are currently used to correct for availability bias in these species (Harker et al., 2022). We therefore logit-transformed the inverse of the proportion of time spent at the surface in puffins (Spencer, 2012), guillemots (Thaxter et al., 2010), razorbills (Thaxter et al., 2010), and red-throated divers (using a value for great northern divers *Gavia immer*; Winiarski et al., 2014). To eliminate divergent transitions that caused a bias in the posterior draws of the puffin model, we removed the December data (n_days_ = 1) and one outlier value (<4 minutes submerged). Whilst we initially ran all models with 4 MCMC chains and with 2,000 iterations per chain, including warmup, we increased the number of iterations of the puffin model to 3,000. We confirmed model convergence via visual inspection of the chains, posterior predictive check plots, and by calculating a Gelman-Rubin convergence statistic 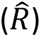 of close to 1 (McElreath, 2020). We used the ‘*marginaleffects*’ R package (Arel-Bundock, 2023) to calculate the marginal effects of each unit change in the explanatory variables. For puffins, guillemots, and razorbills, we used these values to make comparisons between our model estimates of the proportions of daylight hours that the species spent submerged, and the proportion of time spent on the water during which birds were submerged, the quantity used previously within offshore EIAs (Harker et al., 2022). We note that these values are therefore not identical in their derivation, due to us not evaluating flight behaviour here. However, they remain comparable due to the very small proportion of time (typically <2%) that these species spend engaged in flight during the non-breeding season (Duckworth, 2023; Dunn et al., 2020).

We provide values of *pr*(*being visible*), the probability of an animal being at the water’s surface or in flight and therefore available to be recorded on a survey, for the months of the non-breeding season for puffins in the North Sea, guillemots in the North Sea and the UK west coast, razorbills in the North Sea, and red-throated divers that move between the Baltic Sea and the North Sea, those within Icelandic coastal waters, and those off the north coast of Scotland (Table 3). These values were derived by subtracting the posterior distributions of the models’ outputs for the proportion of daylight hours spent submerged (i.e., the mean estimate ± lower and upper 95% confidence intervals) from 1. Values of *pr*(*being visible*) can be used to correct relative abundance estimates of birds by dividing the density of birds observed sitting on the sea and in flight by *pr*(*being visible*) (Barlow et al., 1988). Thus, the values generated by this study can be used in offshore windfarm assessments in a similar way that existing availability bias correction factors are currently used.

**Table 3.**
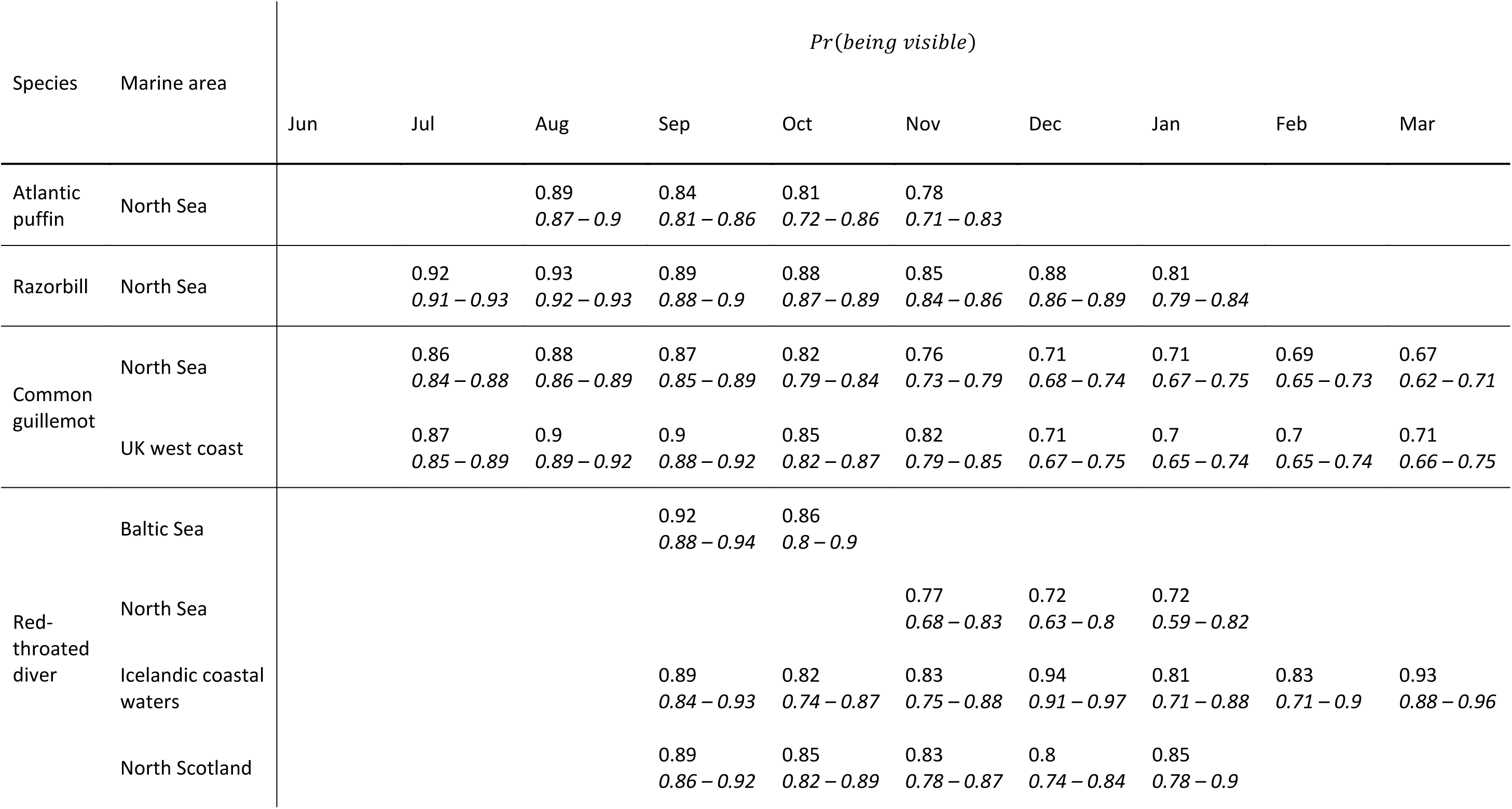
Estimates (and lower and upper 95% confidence intervals in italics) of the probability of being available at the surface or in flight, *pr*(*being visible*), for Atlantic puffins, common guillemots, razorbills, and red-throated divers from breeding sites across northwest Europe during the months of the non-breeding season. These correction factors can be applied to survey estimates by dividing estimates of the density or abundance of birds in flight and on water by *pr*(*being visible*) to give corrected estimates.

| Species | Marine area | *Pr(being visible)* |  |  |  |  |  |  |  |  |  |
| --- | --- | --- | --- | --- | --- | --- | --- | --- | --- | --- | --- |
|  |  | Jun | Jul | Aug | Sep | Oct | Nov | Dec | Jan | Feb | Mar |
| Atlantic puffin | North Sea |  |  | 0.89<br>*0.87 – 0.9* | 0.84<br>*0.81 – 0.86* | 0.81<br>*0.72 – 0.86* | 0.78<br>*0.71 – 0.83* |  |  |  |  |
| Razorbill | North Sea |  | 0.92<br>*0.91 – 0.93* | 0.93<br>*0.92 – 0.93* | 0.89<br>*0.88 – 0.9* | 0.88<br>*0.87 – 0.89* | 0.85<br>*0.84 – 0.86* | 0.88<br>*0.86 – 0.89* | 0.81<br>*0.79 – 0.84* |  |  |
| Common guillemot | North Sea |  | 0.86<br>*0.84 – 0.88* | 0.88<br>*0.86 – 0.89* | 0.87<br>*0.85 – 0.89* | 0.82<br>*0.79 – 0.84* | 0.76<br>*0.73 – 0.79* | 0.71<br>*0.68 – 0.74* | 0.71<br>*0.67 – 0.75* | 0.69<br>*0.65 – 0.73* | 0.67<br>*0.62 – 0.71* |
|  | UK west coast |  | 0.87<br>*0.85 – 0.89* | 0.9<br>*0.89 – 0.92* | 0.9<br>*0.88 – 0.92* | 0.85<br>*0.82 – 0.87* | 0.82<br>*0.79 – 0.85* | 0.71<br>*0.67 – 0.75* | 0.7<br>*0.65 – 0.74* | 0.7<br>*0.65 – 0.74* | 0.71<br>*0.66 – 0.75* |
| Red-throated diver | Baltic Sea |  |  |  | 0.92<br>*0.88 – 0.94* | 0.86<br>*0.8 – 0.9* |  |  |  |  |  |
|  | North Sea |  |  |  |  |  | 0.77<br>*0.68 – 0.83* | 0.72<br>*0.63 – 0.8* | 0.72<br>*0.59 – 0.82* |  |  |
|  | Icelandic coastal waters |  |  |  | 0.89<br>*0.84 – 0.93* | 0.82<br>*0.74 – 0.87* | 0.83<br>*0.75 – 0.88* | 0.94<br>*0.91 – 0.97* | 0.81<br>*0.71 – 0.88* | 0.83<br>*0.71 – 0.9* | 0.93<br>*0.88 – 0.96* |
|  | North Scotland |  |  |  | 0.89<br>*0.86 – 0.92* | 0.85<br>*0.82 – 0.89* | 0.83<br>*0.78 – 0.87* | 0.8<br>*0.74 – 0.84* | 0.85<br>*0.78 – 0.9* |  |  |

## Results

There was temporal variation in the proportion of daylight hours that puffins, guillemots, razorbills, and red-throated divers spent submerged over the months that followed the end of the species’ breeding seasons. Birds spent an average of 1.5 hours submerged each day, this varying between 1% and 59% of daylight hours spent submerged.

The proportion of daylight hours that puffins in the North Sea spent submerged was lowest in July and August (Fig 2A). In July, the confidence intervals encompassed the proportion of time on water submerged previously used to account for availability bias in this species (Spencer, 2012). Although there were wide confidence intervals surrounding estimates of the proportion of daylight hours that puffins spent submerged, this value increased throughout September, October, and November, again overlapping with the previously used value in October before exceeding it in November (Fig 2A).

**Figure 2.**
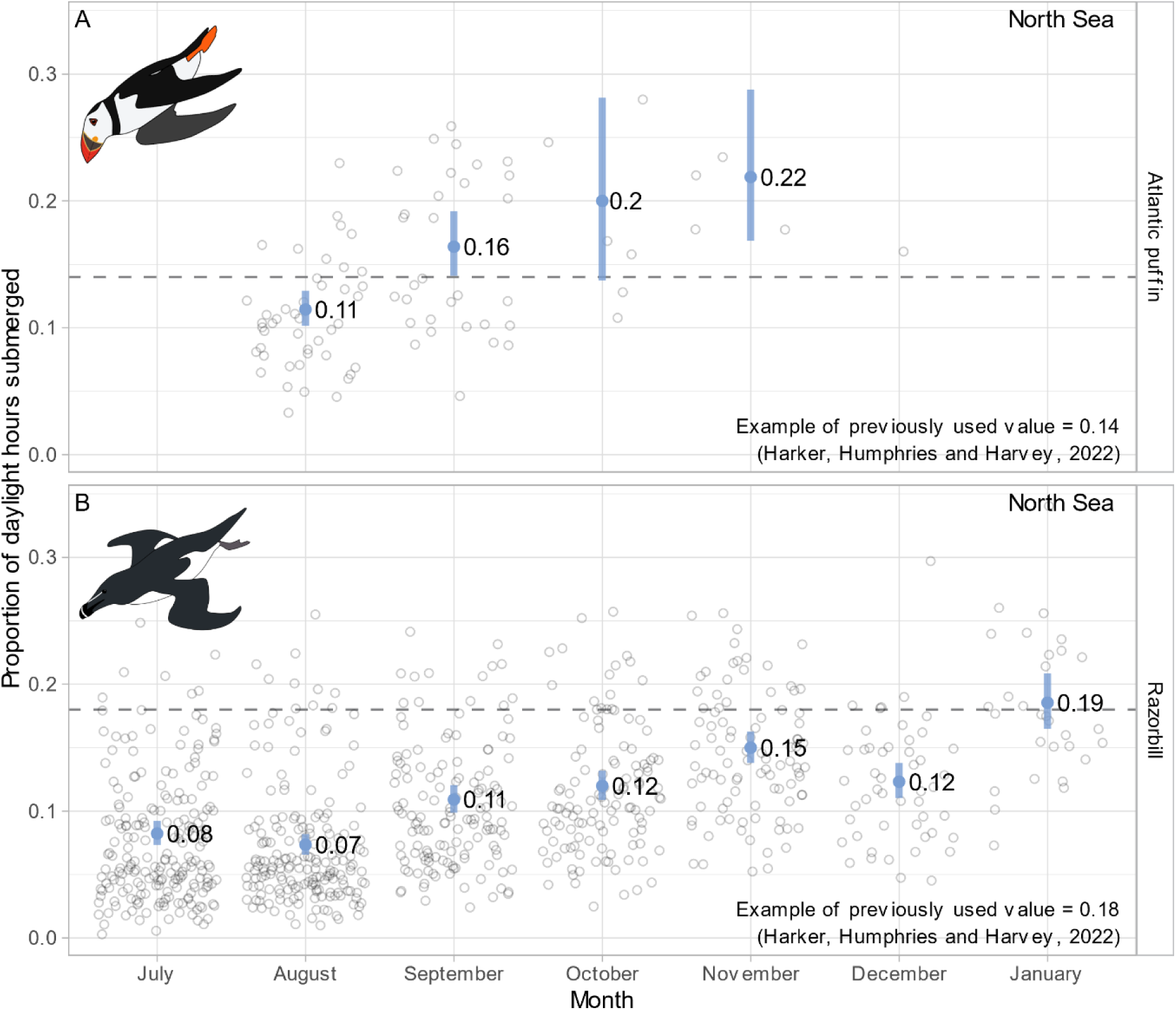
Marginal model predictions of the proportion of daylight hours spent submerged by Atlantic puffins (A) and razorbills (B) wintering within the North Sea, averaged by month, between August and November and July and January for puffins and razorbills, respectively. Blue points illustrate mean marginal predictions and shaded lines illustrate 95% confidence intervals, and the raw data are shown as faint open circles behind. The horizontal dashed lines indicate values of the proportion of time on water submerged previously used during environmental impact assessments for these species (see text for details).

The temporal pattern in the proportion of daylight hours submerged was similar in razorbills wintering around the east coast of the UK; they spent little time submerged below the water’s surface during the daytime periods that immediately followed the breeding season (July – August), but this increased over the course of the winter, only overlapping with a previously used value in January (Thaxter et al., 2010).

The proportion of daylight hours that guillemots spent submerged during their non-breeding season was similar between birds wintering in both the North Sea and off the UK west coast (Fig 3). Initially, following their breeding season, birds spent a relatively low proportion of daylight hours submerged beneath the water’s surface. Indeed, values between July – October were lower than values of the proportion of time on water submerged previously used to correct for availability bias, only exceeding this value in November for birds wintering in the North Sea, and December for birds wintering off the UK west coast (Fig 3). The proportion of daylight hours spent submerged then increased for both groups throughout the sampling period until March (Fig 3).

**Figure 3.**
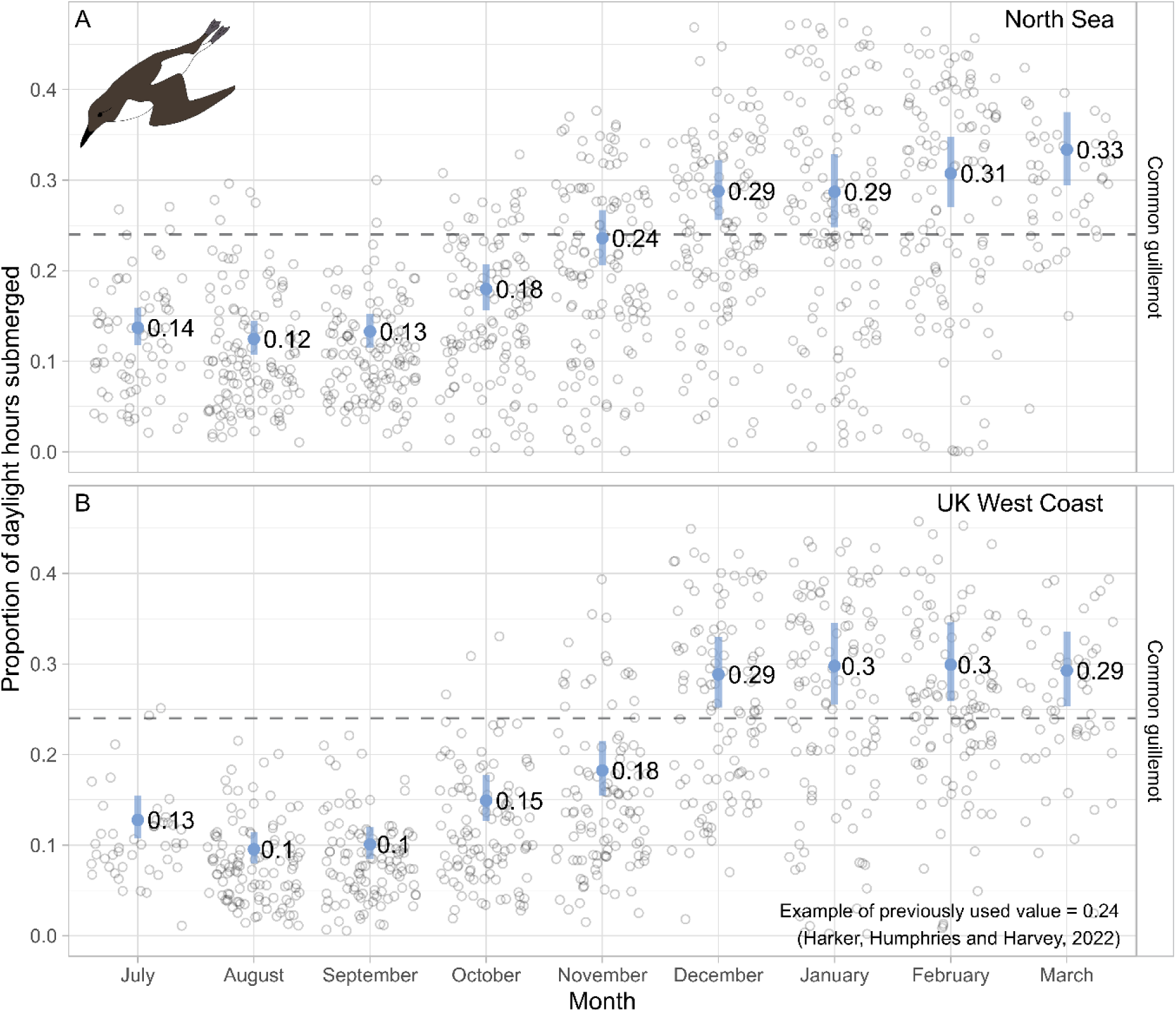
Marginal model predictions of the proportion of daylight hours spent submerged by common guillemots wintering (A) in the North Sea and (B) off the UK west coast, averaged by month, between July and March. Blue points illustrate mean marginal predictions and shaded lines illustrate 95% confidence intervals, and the raw data are shown as faint open circles behind. The horizontal dashed lines indicate a value of the proportion of time on water submerged previously used during environmental impact assessments for this species (see text for details).

Red-throated divers wintering in different areas spent differing proportions of daylight hours submerged. Birds that initially wintered within the Baltic Sea, before moving to the southern North Sea during October, spent an increasing amount of time below the water’s surface following their breeding seasons (Fig. 4A). Although a similar pattern was observed in red-throated divers that wintered around north Scotland, with the proportion of daylight hours submerged increasing between September and December, there was a slight decrease in time spent submerged in January (Fig. 4C). A similar dip in the proportion of daylight hours submerged in December was observed in red-throated divers that wintered in Icelandic coastal waters, with high values being observed in January and February, before another decrease in March (Fig. 4B).

**Figure 4.**
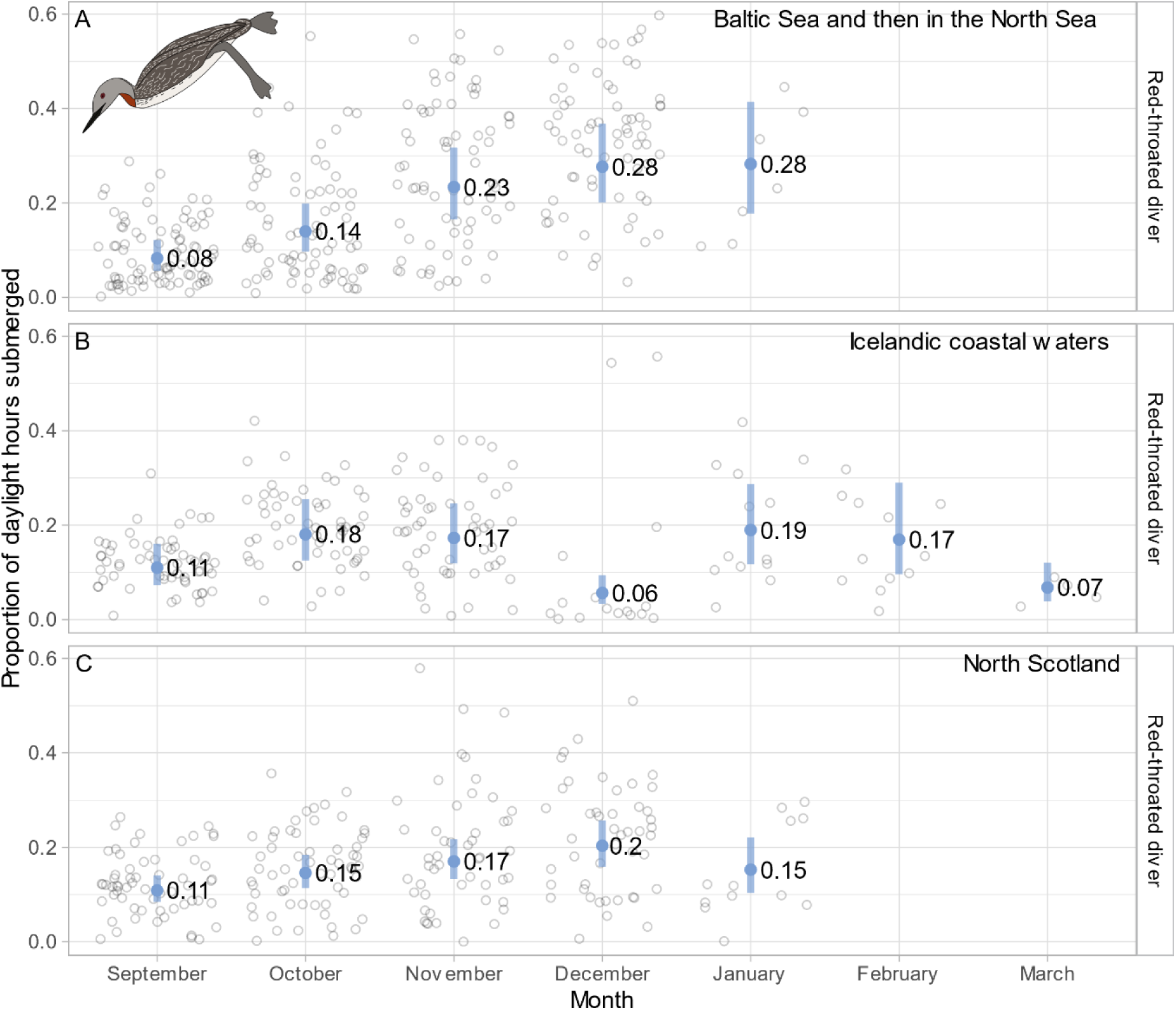
Marginal model predictions of the proportion of daylight hours spent submerged by red-throated divers wintering (A) in the Baltic Sea during September and October and then in the North Sea during November – January, (B) in Icelandic coastal waters, and (C) off north Scotland, averaged by month, between September and March. Blue points illustrate mean marginal predictions and shaded lines illustrate 95% confidence intervals, and the raw data are shown as faint open circles behind.

Temporal, spatial, and interspecific variation in the proportion of daylight hours spent submerged by the puffins, guillemots, razorbills, and red-throated divers led to variation in values of *pr*(*being visible*), the probability of each species being available to be recorded at the water’s surface or in flight during surveys (Table 3). These values vary between 0.67 (95% confidence interval = 0.62 – 0.71) for guillemots wintering in the North Sea in March, and 0.95 for razorbills in a similar area in June (95% confidence interval = 0.93 – 0.96).

## Discussion

There was substantial variation in the proportion of daylight hours spent submerged by puffins, guillemots, razorbills, and red-throated divers in different marine areas throughout their non-breeding seasons. This variation led to differences in the probability of them being available for detection by commonly used survey methods (‘availability bias’, defined as ‘*pr*(*being visible*)’), ultimately influencing estimates of their absolute densities. Our results suggest that previously used correction factors for availability bias (often derived from individuals of different species, or in different areas, or during different seasons) likely provided incorrect estimates of marine bird abundances, the degree of which would have varied by species, time, and area. Indeed, for red-throated divers where no correction factor is regularly used, surveys are currently likely to often yield underestimates of their non-breeding abundances. We therefore encourage the use of correction factors that are estimated from the species, marine area, and month during which the survey is conducted. Our study highlights the importance of using appropriate input data (i.e., data that is as closely related to the surveyed population as possible) when quantifying species abundances and distributions, evidence that is relevant for various applied science studies. We present new values through which to account for variation in availability bias for our four focal species and their corresponding marine areas (Table 3).

Our increased understanding of marine behaviour outside the breeding season is highlighting a mounting number of differences when compared to behaviour during the breeding season (Drummond et al., 2021). In this study, the proportion of daylight hours that all four species spent submerged below the water’s surface varied across the non-breeding season, generally demonstrating an increase over this time (Fig 2-4). This temporal variation is likely to be representative of fluctuations in climatic conditions, and the birds’ subsequent energy requirements and activity budgets (Buckingham et al., 2023; Clairbaux et al., 2021; Fort et al., 2009). Differences in dive behaviour can also be compounded by variation in daylight availability impacting the ability of these visual foragers to gain the energy needed to meet their energetic demands, therefore driving changes in their time activity budgets (Duckworth et al., 2021; Dunn et al., 2020). Indeed, the time that marine birds spend submerged varies over the course of the diurnal cycle (Jardine, 2024), and should be considered within future work. Other life history events such as feather moult (Cherel et al., 2016) and migratory travel can also influence marine bird activity budgets, with migratory populations of red-throated divers (Fig 4A) spending a higher proportion of time submerged than more residential populations (Fig 4B-C). This variation in activity is likely due to a requirement to adjust their behaviour as they moved from one wintering area to another in addition to needing to meet the high energetic costs often associated with travel in marine birds (Dunn et al., 2023).

Here we show that accounting for temporal and spatial variation in behaviour is critical to enhancing our understanding of the consequences of anthropogenic change. In particular, seasonal change in availability bias would strongly affect estimates of seasonal abundance. For example, for common guillemots off the west of Scotland, the correction factor was three times higher in February than in September (Figure 3). It is important that EIA and other assessments do not incorrectly estimate marine bird abundances, something that can occur if inappropriate availability bias values continue to be used. Our results suggest that a hypothetical set of sequential surveys might show a decreasing trend in guillemot abundance if a fixed correction factor was used, but in reality, the trend could be stable if the appropriate varying proportion of time submerged (and hence correction factor) was used. This reinforces the recommendation to use correction factors that acknowledge these temporal trends to garner more accurate estimates of the at-sea abundances of these species outside their breeding seasons.

EIAs for offshore renewable projects typically derive corrected estimates of marine bird abundances and distributions using corrections for availability bias, but here we show that current practices may not be dependable. Currently, raw counts of birds on the water’s surface are divided by *pr*(*being visible*), therefore increasing the estimate above the original count, before adding the number of flying birds. For example, if 3,000 common guillemots were recorded on a digital aerial survey in the North Sea during September, previously used correction factors would have estimated that there were 3,947 birds, with no associated availability bias error. Instead, our species-, time-, and area-specific correction factor (outlined in Table 3) would have produced an estimate of the number of birds in the area being 3,448 individuals (95% confidence interval = 3,370 – 3,529). Furthermore, if in January the survey recorded 3,000 common guillemots, 10% of which were in flight, previously used correction factors would have estimated 3,938 birds, with no associated availability bias error. Contrastingly, our correction factor (Table 3) would instead suggest that the number of birds likely in the area was 4,225 individuals (95% confidence interval = 4,000 – 4,478). Furthermore, in red-throated divers where no correction factors are currently regularly used, population estimates may be missing almost one third of the red-throated divers in the North Sea during December and January (Figure 4A). These examples demonstrate that previous estimates of the abundances and distributions of these marine birds during winter may well have been incorrect, as they fall outside the confidence intervals generated via this new approach, with implications for assessing the potential consequences of anthropogenic activities such as offshore windfarm developments.

Our study shows how incorporating our increasing understanding of temporal and spatial variation in the behaviour of non-breeding marine birds has consequences for the assessment of anthropogenic activities within the marine environment. The data presented here are most relevant to EIA within the context of offshore wind development but would also have relevance for protected area designation and the investigation of long-term change across marine systems. We encourage those making assessments elsewhere to use and/or collect the most appropriate information to their surveyed areas or studied populations, as was originally conceived in attempts to correct for availability bias (Barlow et al., 1988). In doing so, we recommend the use of larger sample sizes, longer term logger deployments, and improved non-breeding area estimates to enhance future insights, something that is becoming increasingly tangible with the continued modernisation and miniaturisation of biologging devices. As these developments continue, it will be important to also consider longer-term interannual temporal variation, and the movements of juvenile and immature individuals (Fayet et al., 2024; Merkel & Strøm, 2023). All of this is particularly timely during a period when marine birds are suffering catastrophic declines (Dias et al., 2019) in tandem with our oceans becoming increasingly intensively used for renewable energy developments, oil and gas drilling, transport, and sand extraction (Halpern et al., 2019). Ultimately, when seeking to monitor species abundances and distributions we must collect, harness, and appropriately interpret relevant datasets, embracing the nuances and seeking to understand the uncertainty of these data and the underlying ecological processes (Chadwick et al., 2023).

## Authors’ contributions

Sue O’Brien, Jonathan Green, and James Duckworth conceived the ideas; Ruth Dunn, James Duckworth and Lila Buckingham managed the data; Ruth Dunn analysed the data; Ruth Dunn led the data visualisation with marine bird illustrations provided by Lila Buckingham; Ruth Dunn led the writing of the manuscript with support from Jonathan Green. All authors contributed critically to the drafts.

## Acknowledgements

We extend our thanks to all individuals and groups who contributed to data collection, namely David Jardine (Colonsay auks), Mark Newell, Sarah Wanless, Mike Harris, and Sophie Bennett (Isle of May auks), Robin Ward and Chris Heward (Treshnish Isles Auk Ringing Group; auks), Ewan Weston (Whinnyfold auks), Jim Williams and Stuart Williams (Orkney red-throated divers), David Okill and Logan Johnson (Shetland red-throated divers), Aevar Petersen (Iceland red-throated divers), and Roni Vaisainen and Pepe Lehikoinen (Finland red-throated divers). Atlantic puffin and razorbill fieldwork was funded by NERC (NERC National Capability, NERC/Department for Environment, Food and Rural Affairs Marine Ecosystems Research Programme Ref NE/L003082/1), the Royal Society for the Prevention of Cruelty to Animals and Equinor, and guillemot fieldwork was funded by Vattenfall, Marine Scotland Science, Equinor (as part of Hywind Scotland’s Environmental Monitoring Programme) and SEATRACK. Red-throated diver fieldwork was managed by JNCC and funded by The Crown Estate, Ørsted, Equinor, Vattenfall, and Hartley Anderson Ltd (BEIS Offshore Energy SEA Research Fund). We extend thanks to the Ministry for the Environment and Natural Resources in Iceland for support during the study in Iceland, and NatureScot for access to the Isle of May. We are also grateful to all those involved with giving permission to undertake this work including the Centre for Economic Development, Transport and the Environment in Finland, the National Parks, Finland, the British Trust for Ornithology, Scottish Natural Heritage, the Royal Society for the Protection of Birds. RED was funded by the Bertarelli Foundation as part of the Bertarelli Programme in Marine Science. JAD’s PhD studentship was funded by a Natural Environment Research Council (NERC) CASE PhD studentship in collaboration with JNCC, as part of the ACCE Doctoral Training Partnership. The authors would also like to extend their thanks to Tim Fay and Emma Ahart and their support for this work, along with members of the Project Steering Group they assembled, all of whom provided insight that fed into the direction of the analyses and the final manuscript. We thank Dr Michael Roast and an anonymous reviewer for helpful comments on an earlier draft of this manuscript.

All applicable institutional and/or national guidelines for the care and use of animals were followed.

## Data availability statement

No new data were collected for the study.

## Data sources

Atlantic puffin and razorbill dive data: Dunn R.E., Wanless S., Green J.A., Harris, M.P., Daunt F. 2019. Dive times and depths of auks (Atlantic puffin, common guillemot and razorbill) from the Isle of May outside the breeding season. Environmental Information Data Centre, https://doi.org/10.5285/6ab0ee70-96f8-41e6-a3e3-6f4c31fa5372.

Guillemot dive data: Buckingham, L., Daunt, F., Bogdanova, M.I., Furness, R.W., Bennett, S., Duckworth, J., Dunn, R.E., Wanless, S., Harris, M.P., Jardine, D.C., Newell, M.A., Ward, R.M., Weston, E.D. and Green, J.A. 2022. Energetic synchrony throughout the non-breeding season in common guillemots from four colonies. Zenodo, https://zenodo.org/record/7327472#.Y7_FOHbMK4s.

Red-throated diver dive data are detailed in Duckworth, et al. (2021). Data will be held at the public JNCC repository at https://hub.jncc.gov.uk/ email enquiries can be sent to.

## References

Arel-Bundock, V. (2023). *marginaleffect*s: Predictions, comparisons, *slopes, marginal means, and hypothesis tests*. R package version 0.9. https://marginaleffects.com/

Barlow, J., Oliver, C. W., Jackson, T. D., & Taylor, B. L. (1988). Harbor porpoise, Phocoena phocoena, abundance estimation for California, Oregon, and Washington: II. Aerial surveys. Fishery Bulletin, 86, 433–444.

Boivin, N. L., Zeder, M. A., Fuller, D. Q., Crowther, A., Larson, G., Erlandson, J. M., Denham, T., & Petraglia, M. D. (2016). Ecological consequences of human niche construction: Examining long-term anthropogenic shaping of global species distributions. Proceedings of the National Academy of Sciences of the United States of America, 113(23), 6388–6396.

Buckingham, L., Bogdanova, M. I., Green, J. A., Dunn, R. E., Wanless, S., Bennett, S., Bevan, R. M., Call, A., Canham, M., Corse, C. J., Harris, M. P., Heward, C. J., Jardine, D. C., Lennon, J., Parnaby, D., Redfern, C. P. F., Scott, L., Swann, R. L., Ward, R. M., … Daunt, F. (2022). Interspecific variation in non-breeding aggregation: a multi-colony tracking study of two sympatric seabirds. Marine Ecology Progress Series, 684, 181–197.

Buckingham, L., Daunt, F., Bogdanova, M. I., Furness, R. W., Bennett, S., Duckworth, J., Dunn, R. E., Wanless, S., Harris, M. P., Jardine, D. C., Newell, M. A., Ward, R. M., Weston, E. D., & Green, J. A. (2023). Energetic synchrony throughout the non-breeding season in common guillemots from four colonies. Journal of Avian Biology, 2023, e03018.

Buckland, S. T., Burt, M. L., Rexstad, E. A., Mellor, M., Williams, A. E., & Woodward, R. (2012). Aerial surveys of seabirds: the advent of digital methods. The Journal of Applied Ecology, 49(4), 960– 967.

Bürkner, P.-C. (2017). brms: An R Package for Bayesian Multilevel Models Using Stan. Journal of Satistical Software, 80, 1–28.

Certain, G., & Bretagnolle, V. (2008). Monitoring seabirds population in marine ecosystem: The use of strip-transect aerial surveys. Remote Sensing of Environment, 112(8), 3314–3322.

Chadwick, F. J., Haydon, D. T., Husmeier, D., Ovaskainen, O., & Matthiopoulos, J. (2023). LIES of omission: complex observation processes in ecology. Trends in Ecology & Evolution. 10.1016/j.tree.2023.10.009

Cherel, Y., Quillfeldt, P., Delord, K., & Weimerskirch, H. (2016). Combination of at-sea activity, geolocation and feather stable isotopes documents where and when seabirds molt. Frontiers in Ecology and Evolution, 4, 3.

Clairbaux, M., Mathewson, P., Porter, W., Fort, J., Strøm, H., Moe, B., Fauchald, P., Descamps, S., Helgason, H. H., Bråthen, V. S., Merkel, B., Anker-Nilssen, T., Bringsvor, I. S., Chastel, O., Christensen-Dalsgaard, S., Danielsen, J., Daunt, F., Dehnhard, N., Erikstad, K. E., … Grémillet, D. (2021). North Atlantic winter cyclones starve seabirds. Current Biology, 31(17), 3964–3971.

Delong, L., Shuyao, W., Laibao, L., Yatong, Z., & Shuangcheng, L. (2018). Vulnerability of the global terrestrial ecosystems to climate change. Global Change Biology, 24(9), 4095–4106.

Dias, M. P., Martin, R., Pearmain, E. J., Burfield, I. J., Small, C., Phillips, R. A., Yates, O., Lascelles, B., Borboroglu, P. G., & Croxall, J. P. (2019). Threats to seabirds: A global assessment. Biological Conservation, 237, 525–537.

Dierschke, V., Furness, R. W., & Garthe, S. (2016). Seabirds and offshore wind farms in European waters: Avoidance and attraction. Biological Conservation, 202, 59–68.

Drummond, B. A., Orben, R. A., Christ, A. M., Fleishman, A. B., Renner, H. M., Rojek, N. A., & Romano, M. D. (2021). Comparing non-breeding distribution and behavior of red-legged kittiwakes from two geographically distant colonies. PloS One, 16(7), e0254686.

Duckworth, J. (2023). Using behavioural and energetic insights to assess the impacts of displacement from offshore wind farms on red-throated divers (Gavia stellata) [PhD, University of Liverpool]. 10.17638/03170909

Duckworth, J., O’Brien, S., Petersen, I. K., Petersen, A., Benediktsson, G., Johnson, L., Lehikoinen, P., Okill, D., Väisänen, R., Williams, J., Williams, S., Daunt, F., & Green, J. A. (2021). Spatial and temporal variation in foraging of breeding red-throated divers. Journal of Avian Biology, 52(6), e02702.

Duckworth, J., O’Brien, S., Petersen, I. K., Petersen, A., Benediktsson, G., Johnson, L., Lehikoinen, P., Okill, D., Väisänen, R., Williams, J., Williams, S., Daunt, F., & Green, J. A. (2022). Winter locations of red-throated divers from geolocation and feather isotope signatures. Ecology and Evolution, 12(8), e9209.

Duckworth, J., O’Brien, S., Väisänen, R., Lehikoinen, P., Petersen, I. K., Daunt, F., & Green, J. A. (2020). First biologging record of a foraging red-throated loon gavia stellata shows shallow and efficient diving in freshwater environments. Marine Ornithology, 48, 17–22.

Dunn, R. E., Duckworth, J., & Green, J. A. (2023). A framework to unlock marine bird energetics. The Journal of Experimental Biology, 226(24), jeb246754.

Dunn, R. E., Wanless, S., Daunt, F., Harris, M. P., & Green, J. A. (2020). A year in the life of a north Atlantic seabird: behavioural and energetic adjustments during the annual cycle. Scientific Reports, 10, 5993.

Dunn, R. E., Wanless, S., Green, J. A., Harris, M. P., & Daunt, F. (2019). Effects of body size, sex, parental care and moult strategies on auk diving behaviour outside the breeding season. Journal of Avian Biology, 50(7), 1–14.

Edgar, G. J., Stuart-Smith, R. D., Willis, T. J., Kininmonth, S., Baker, S. C., Banks, S., Barrett, N. S., Becerro, M. A., Bernard, A. T. F., Berkhout, J., Buxton, C. D., Campbell, S. J., Cooper, A. T., Davey, M., Edgar, S. C., Försterra, G., Galván, D. E., Irigoyen, A. J., Kushner, D. J., … Thomson, R. J. (2014). Global conservation outcomes depend on marine protected areas with five key features. Nature, 506(7487), 216–220.

Eriksson, M. O. G., Blomqvist, D., Hake, M., & Johansson, O. C. (1990). Parental feeding in the Red- throated Diver *Gavia stellata*. Ibis, 132, 1–13.

Fayet, A. L., Shoji, A., & Guilford, T. (2024). Post-fledging movements of Atlantic Puffins from Skomer Island. Seabird, 36. 10.61350/sbj.36.1

Fort, J., Porter, W. P., & Grémillet, D. (2009). Thermodynamic modelling predicts energetic bottleneck for seabirds wintering in the northwest Atlantic. The Journal of Experimental Biology, 212, 2483–2490.

Guisan, A., Tingley, R., Baumgartner, J. B., Naujokaitis-Lewis, I., Sutcliffe, P. R., Tulloch, A. I. T., Regan, T. J., Brotons, L., McDonald-Madden, E., Mantyka-Pringle, C., Martin, T. G., Rhodes, J. R., Maggini, R., Setterfield, S. A., Elith, J., Schwartz, M. W., Wintle, B. A., Broennimann, O., Austin, M., … Buckley, Y. M. (2013). Predicting species distributions for conservation decisions. Ecology Letters, 16(12), 1424–1435.

Halpern, B. S., Frazier, M., Afflerbach, J., Lowndes, J. S., Micheli, F., O’Hara, C., Scarborough, C., & Selkoe, K. A. (2019). Recent pace of change in human impact on the world’s ocean. Scientific Reports, 9(1), 11609.

Halpern, B. S., Frazier, M., Potapenko, J., Casey, K. S., Koenig, K., Longo, C., Lowndes, J. S., Rockwood, R. C., Selig, E. R., Selkoe, K. A., & Walbridge, S. (2015). Spatial and temporal changes in cumulative human impacts on the world’s ocean. Nature Communications, 6(1), 7615.

Harker, J., Humphries, G., & Harvey, J. (2022). Berwick Bank Wind Farm Offshore Environmental Impact Assessment. HiDef Aerial Surveying Ltd.

Harris, M. P., Daunt, F., Newell, M., Phillips, R. A., & Wanless, S. (2010). Wintering areas of adult Atlantic puffins *Fratercula arctica* from a North Sea colony as revealed by geolocation technology. Marine Biology, 157(4), 827–836.

Harris, M. P., & Wanless, S. (2011). The Puffin. T & A D Poyser.

Heiss, A. (2021, November 8). A guide to modeling proportions with Bayesian beta and zero-inflated beta regression models. 10.59350/7p1a4-0tw75

Henson, S. A., Beaulieu, C., Ilyina, T., John, J. G., Long, M., Séférian, R., Tjiputra, J., & Sarmiento, J. L. (2017). Rapid emergence of climate change in environmental drivers of marine ecosystems. Nature Communications, 8(1), 14682.

Irwin, C., Scott, M. S., Humphries, G., & Webb, A. (2019). *HiDef report to Natural England - Digital video aerial surveys of red-throated diver in the Outer Thames Estuary Special Protection Area* 2018 (No. 260). Natural England Commissioned Reports.

Jardine, D. C. (2024). Feeding ecology of wintering Great Northern Divers *Gavia immer* in Argyll, Scotland. Seabird, 36. 10.61350/sbj.36.4

Jetz, W., McGeoch, M. A., Guralnick, R., Ferrier, S., Beck, J., Costello, M. J., Fernandez, M., Geller, G. N., Keil, P., Merow, C., Meyer, C., Muller-Karger, F. E., Pereira, H. M., Regan, E. C., Schmeller, D. S., & Turak, E. (2019). Essential biodiversity variables for mapping and monitoring species populations. Nature Ecology & Evolution, 3(4), 539–551.

Kleinschmidt, B., Burger, C., Dorsch, M., Nehls, G., Heinänen, S., Morkūnas, J., Žydelis, R., Moorhouse-Gann, R. J., Hipperson, H., Symondson, W. O. C., & Quillfeldt, P. (2019). The diet of red-throated divers (*Gavia stellata*) overwintering in the German Bight (North Sea) analysed using molecular diagnostics. Marine Biology, 166(6), 77.

Lescure, L., Gulka, J., & Davoren, G. K. (2023). Increased Foraging Effort and Reduced Chick Condition of Razorbills under Lower Prey Biomass in Coastal Newfoundland. *Canada.”* Marine Ecology Progress Series, 709, 109–123.

Linnebjerg, J. F., Fort, J., Guilford, T., Reuleaux, A., Mosbech, A., & Frederiksen, M. (2013). Sympatric breeding auks shift between dietary and spatial resource partitioning across the annual cycle. PloS One, 8(8), 1–10.

McElreath, R. (2020). Statistical Rethinking: A Bayesian Course with Examples in R and Stan (2nd ed., pp. 1–612). Chapman & Hall.

Merkel, B., & Strøm, H. (2023). Post-colony swimming migration in the genus *Uria*. Journal of Avian Biology, 2024(1–2), e03153.

NatureScot. (2023a, January). Guidance Note 1: Guidance to support Offshore Wind Applications: Marine Ornithology - Overview. NatureScot. https://www.nature.scot/doc/guidance-note-1-guidance-support-offshore-wind-applications-marine-ornithology-overview

NatureScot. (2023b, January). Offshore Wind Ornithological Impact Assessment - Review of Digital Aerial Survey Methods. NatureScot. https://www.nature.scot/doc/offshore-wind-ornithological-impact-assessment-review-digital-aerial-survey-methods

Phillips, R. A., Lewis, S., González-Solís, J., & Daunt, F. (2017). Causes and consequences of individual variability and specialization in foraging and migration strategies of seabirds. Marine Ecology Progress Series, 578, 117–150.

R Core Team. (2022). *R: A language and environment for statistical computing* (4.3.2.). R Foundation for Statistical Computing. http://www.r-project.org/

Sergio, F., Newton, I., Marchesi, L., & Pedrini, P. (2006). Ecologically justified charisma: preservation of top predators delivers biodiversity conservation. The Journal of Applied Ecology, 43(6), 1049–1055.

Shoji, A., Elliott, K. H., Fayet, A. L., Boyle, D., Perrins, C., & Guilford, T. (2015). Foraging behaviour of sympatric razorbills and puffins. Marine Ecology Progress Series, 520, 257–267.

Soares-Ramos, E. P. P., de Oliveira-Assis, L., Sarrias-Mena, R., & Fernández-Ramírez, L. M. (2020). Current status and future trends of offshore wind power in Europe. Energy, 202, 117787.

Spencer, S. M. (2012). Diving Behavior and Identification of Sex of Breeding Atlantic Puffins (Fratercula arctica), and Nest-Site Characteristics of Alcids on Petit Manan Island, Maine [Masters, University of Massachusetts Amherst]. 10.7275/2740959

Thaxter, C. B., & Burton, N. H. K. (2009). High definition imagery for surveying seabirds and marine mammals: a review of recent trials and development of protocols. COWRIE BTO Wshop-09 report to COWRIE Ltd., London.

Thaxter, C. B., Wanless, S., Daunt, F., Harris, M. P., Benvenuti, S., Watanuki, Y., Grémillet, D., & Hamer, K. C. (2010). Influence of wing loading on the trade-off between pursuit-diving and flight in common guillemots and razorbills. The Journal of Experimental Biology, 213, 1018–1025.

Wanless, S., Albon, S. D., Daunt, F., Sarzo, B., Newell, M. A., Gunn, C., Speakman, J. R., & Harris, M. P. (2023). Increased parental effort fails to buffer the cascading effects of warmer seas on common guillemot demographic rates. The Journal of Animal Ecology, 92(8), 1622–1638.

Winiarski, K. J., Burt, M. L., Rexstad, E., Miller, D. L., Trocki, C. L., Paton, P. W. C., & McWilliams, S. R. (2014). Integrating aerial and ship surveys of marine birds into a combined density surface model: A case study of wintering Common Loons. The Condor, 116(2), 149–161.

