## Supporting Information S1 and S2 for "Temporal and spatial variability in availability bias has consequences for marine bird abundance estimates during the non-breeding season"

S1. Estimating daylength across non-breeding distributions

To categorise dive incidences as occurring during either daylight, twilight, or nighttime, we extracted population-specific area estimates throughout the non-breeding season. To do this, we obtained centroids of each population’s monthly non-breeding distribution. For puffins from the Isle of May, we extracted a single centroid from the kernel density distributions of their non-breeding distributions (August – December) presented within Figure 1 of (Harris *et al.* 2010). For guillemots, we extracted centroids from monthly the 50% kernel density contours presented in Figure 1 of (Buckingham *et al.* 2023). For razorbills, we extracted centroids from the post-breeding moult (August – September) and midwinter (December – January) 50% kernel density contours presented in Figures 2 and 3 of (Buckingham *et al.* 2022). For red-throated divers, we extracted centroids from the early winter (September – December) 50% kernel density distributions presented in Figures 1–3 of (Duckworth *et al.* 2022). Red-throated divers breeding in Finland migrated from the Baltic Sea to the North Sea during October and November, and so we used two areas for these birds: a Baltic Sea area during September and October, and a North Sea area during November and December. We adjusted the Baltic Sea area to be further south of the 50% kernel density distribution, thereby correcting for the tag being shaded by leg tucking which produced a northern bias (Duckworth *et al.* 2022).

Whilst the areas of the individuals equipped with time-depth-recorders in this study may be slightly different from those used to generate the extent of daylight (sun was >0 degrees above the horizon) and twilight (sun was 0–15 degrees below the horizon) used in this study, we use a liberal definition of daylight and twilight to include as much sub-surface behaviour as possible. Furthermore, the changes in the duration of daylight and twilight periods across each of the population’s ranges are < 20 minutes for most of the study period.

S2. Sensitivity analysis of location estimates

Table 3. Estimates (and lower and upper 95% confidence intervals in italics) of the probability of being available at the surface or in flight, $Pr\left( being visible \right)$, for common guillemots from breeding sites across northwest Europe during the months of the non-breeding season, based on the centroids of their wintering areas (North Sea and UK west coast), as well as the lower and upper standard deviation values of all the locations for each population in each month, demonstrating that differences are negligible.

| Species | Marine area | $Pr\left( being visible \right)$ | | | | | | | | |
| --- | --- | --- | --- | --- | --- | --- | --- | --- | --- | --- |
|  |  | Jul | Aug | Sep | Oct | Nov | Dec | Jan | Feb | Mar |
| Wintering area centroids | North Sea | 0.86  *0.84 – 0.88* | 0.88  *0.86 – 0.89* | 0.87  *0.85 – 0.89* | 0.82  *0.79 – 0.84* | 0.76  *0.73 – 0.79* | 0.71  *0.68 – 0.74* | 0.71  *0.67 – 0.75* | 0.69  *0.65 – 0.73* | 0.67  *0.62 – 0.71* |
|  | UK west coast | 0.87  *0.85 – 0.89* | 0.9  *0.89 – 0.92* | 0.9  *0.88 – 0.92* | 0.85  *0.82 – 0.87* | 0.82  *0.79 – 0.85* | 0.71  *0.67 – 0.75* | 0.7  *0.65 – 0.74* | 0.7  *0.65 – 0.74* | 0.71  *0.66 – 0.75* |
| Mean minus the standard deviation of the wintering area latitudes | North Sea | 0.86  *0.84 – 0.88* | 0.88  *0.86 – 0.89* | 0.87  *0.85 – 0.89* | 0.82  *0.79 – 0.85* | 0.77  *0.73 – 0.8* | 0.72  *0.68 – 0.75* | 0.72  *0.68 – 0.75* | 0.7  *0.65 – 0.74* | 0.67  *0.63 – 0.71* |
|  | UK west coast | 0.87  *0.85 – 0.9* | 0.91  *0.89 – 0.92* | 0.9  *0.88 – 0.92* | 0.85  *0.82 – 0.87* | 0.82  *0.79 – 0.85* | 0.71  *0.67 – 0.75* | 0.71  *0.66 – 0.75* | 0.7  *0.66 – 0.75* | 0.71  *0.66 – 0.75* |
| Mean plus the standard deviation of the wintering area latitudes | North Sea | 0.86  *0.84 – 0.88* | 0.88  *0.86 – 0.89* | 0.87  *0.85 – 0.88* | 0.82  *0.79 – 0.84* | 0.76  *0.73 – 0.79* | 0.71  *0.67 – 0.74* | 0.71  *0.67 – 0.75* | 0.69  *0.65 – 0.73* | 0.66  *0.62 – 0.7* |
|  | UK west coast | 0.87  *0.85 – 0.89* | 0.9  *0.88 – 0.92* | 0.9  *0.88 – 0.91* | 0.85  *0.82 – 0.87* | 0.82  *0.78 – 0.84* | 0.71  *0.67 – 0.75* | 0.7  *0.65 – 0.74* | 0.7  *0.65 – 0.74* | 0.71  *0.66 – 0.75* |

References

Buckingham, L., Bogdanova, M.I., Green, J.A., Dunn, R.E., Wanless, S., Bennett, S., Bevan, R.M., Call, A., Canham, M., Corse, C.J., Harris, M.P., Heward, C.J., Jardine, D.C., Lennon, J., Parnaby, D., Redfern, C.P.F., Scott, L., Swann, R.L., Ward, R.M., Weston, E.D., Furness, R.W. & Daunt, F. (2022) Interspecific variation in non-breeding aggregation: a multi-colony tracking study of two sympatric seabirds. *Marine Ecology Progress Series*, **684**, 181–197.

Buckingham, L., Daunt, F., Bogdanova, M.I., Furness, R.W., Bennett, S., Duckworth, J., Dunn, R.E., Wanless, S., Harris, M.P., Jardine, D.C., Newell, M.A., Ward, R.M., Weston, E.D. & Green, J.A. (2023) Energetic synchrony throughout the non-breeding season in common guillemots from four colonies. *Journal of Avian Biology*, **2023**, e03018.

Duckworth, J., O’Brien, S., Petersen, I.K., Petersen, A., Benediktsson, G., Johnson, L., Lehikoinen, P., Okill, D., Väisänen, R., Williams, J., Williams, S., Daunt, F. & Green, J.A. (2022) Winter locations of red-throated divers from geolocation and feather isotope signatures. *Ecology and Evolution*, **12**, e9209.

Harris, M.P., Daunt, F., Newell, M., Phillips, R.A. & Wanless, S. (2010) Wintering areas of adult Atlantic puffins *Fratercula arctica* from a North Sea colony as revealed by geolocation technology. *Marine Biology*, **157**, 827–836.
